# The effect of device configuration and patient’s body composition on image artifact and RF heating of deep brain stimulation devices during MRI at 1.5T and 3T

**DOI:** 10.1101/2020.04.09.035030

**Authors:** Bhumi Bhusal, Bach T. Nguyen, Jasmine Vu, Behzad Elahi, Joshua Rosenow, Mark J. Nolt, Roberto Lopez-Rosado, Julie Pilitsis, Marisa DiMarzio, Laleh Golestanirad

## Abstract

**BACKGROUND:** Patients with deep brain stimulation (DBS) implants have limited access to MRI due to safety concerns associated with RF-induced heating. Currently, MRI in these patients is allowed only in 1.5T horizontal scanners and with pulse sequences with reduced power. Nevertheless, off-label use of MRI at 3T is increasingly reported based on limited safety assessments. Here we present results of systematic RF heating measurements for two commercially available DBS systems during MRI at 1.5T and 3T.

**PURPOSE:** To assess the effect of imaging landmark, DBS lead configuration, and patient body composition on RF heating of DBS leads during MRI at 1.5 T and 3T.

**STUDY TYPE:** Phantom study.

**POPULATION/SUBJECTS/PHANTOM/SPECIMEN/ANIMAL MODEL:** Gel phantoms and cadaver brain.

**FIELD STRENGTH/SEQUENCE:** 1.5T and 3T, T1-weighted turbo spin echo.

**ASSESSMENT:** RF heating was measured at tips of DBS leads implanted in brain-mimicking gel.

**STATISTICAL TESTS:** None.

**RESULTS:** We observed substantial fluctuation in RF heating mainly affected by phantom composition and DBS lead configuration, ranging from 0.14°C to 23.73°C at 1.5 T, and from 0.10°C to 7.39°C at 3T. The presence of subcutaneous fat substantially altered RF heating at electrode tips (−3.06°C < Δ*T* < 19.05°C). Introducing concentric loops in the extracranial portion of the lead at the surgical burr hole reduced RF heating by up to 89% at 1.5T and up to 98% at 3T compared to worst case heating scenarios.

**DATA CONCLUSION:** Device configuration and patient body composition significantly altered the RF heating of DBS leads during MRI at 1.5T and 3T. Interestingly, certain lead trajectories consistently reduced RF heating and image artifact over different imaging landmarks, RF frequencies, and phantom compositions. Such trajectories could be implemented in patients with minimal disruption to the surgical workflow.

## Introduction

Deep brain stimulation (DBS) is a well-established treatment for severe movement disorders such as drug-resistant Parkinson’s disease (1,2), with rapidly growing emerging applications in psychiatric and affective disorders (3,4). DBS uses an implantable pulse generator (IPG) in the chest to send electric pulses to electrodes implanted in deep brain areas via subcutaneous leads and extensions to modulate the neuronal behavior. Patients with DBS devices can highly benefit from MRI, both for target verification, and for post-operative monitoring of treatment-induced changes in the brain function. Unfortunately, however, postoperative application of MRI is limited for these patients due to risks of radio frequency (RF) heating of the tissue surrounding electrode contacts, a phenomenon known as the “antenna effect” (5–8). Although MR-conditional DBS devices are available, conditions of imaging are restrictive: only 1.5T horizontal scanners are allowed, and only pulse sequences with reduced power (global SAR ≤ 0.1W/kg or B1rms ≤ 2 μT) are recommended (9–11). Complying with manufacturer’s guidelines is difficult in practice, as MRI protocols for optimal visualization of DBS target structures tend to have much higher SAR than what current guidelines allow. Additionally, there is higher contrast to noise ratio at higher fields (e.g., 3T) which significantly helps differentiating small adjacent DBS targets. Consequently, off-label use of MRI for DBS imaging is increasingly reported, both at higher static fields (e.g., 3T), and with higher SAR/B1 levels at 1.5T. Such off-label studies have relied on in-home safety assessments mostly performed in ASTM-like Phantoms (i.e., the box shaped phantom filled with a single material), and with limited number of DBS device configurations (12,13). This oversimplified approach to MRI safety assessment is concerning for two reasons. First, as RF heating is highly sensitive to implant’s configuration and its orientation with respect to MRI electric field (14–22), and as trajectory of DBS leads is substantially variant among patients (23), basing safety inferences on few tested configurations can expose a large number of patients to undue risk. Second, conclusions drawn from studies in single-material box-shaped phantoms may not apply to the realistic cases, as the distribution of electromagnetic fields depend upon the geometry and composition of the sample. A recent study which reported a10-fold discrepancy in the calculated SAR around the implants tested in ASTM phantoms versus human body models underscores this concern (24). Therefore, studies that are more comprehensive and account for different implant configurations, imaging landmarks, and phantom compositions are required to add confidence on the safety of wider use of MRI in patients with DBS implants.

In this work, we report results of a systematic study of effect of device configuration, patient’s body composition, and imaging landmark on RF heating of two commercially available DBS devices from different manufacturers, during MRI at 1.5T and 3T. We designed and created anthropomorphic phantoms consisting of 3D-printed skulls filled with brain mimicking gel and human-shaped head and torso containers filled either with saline solution, or a combination of saline and oil on top representing subcutaneous fat. A total of 192 RF heating measurement were performed assessing the effect of phantom composition, lead configuration, imaging landmark and RF frequency on DBS RF heating during MRI. We found that cases with added oil layer which represented patients with more subcutaneous fat produced substantially higher heating in some cases, a consequential observation which has not been reported before. Similarly, variation in extra-cranial trajectories of DBS leads significantly altered RF heating at DBS electrode contacts within the brain. Interestingly, some lead trajectories consistently reduced RF heating over different body compositions, RF transmit frequencies, and imaging landmarks. Namely, introducing concentric loops at the location of surgical burr hole reduced RF heating for the devices by 89% and 88% at 1.5T, and by 98% and 87% at 3T during head imaging compared to the corresponding worst heating scenarios.

Besides tissue heating, the image artifact around DBS electrodes due to the distortion of B1 field by induced RF currents in the lead is another issue affecting clinical usefulness of images. Specifically, in the case of long conductive wires such as DBS leads, RF artifact can dominate the susceptibility effect (25) and thus, severely hinder electrode localization. We hypothesize that lead trajectories that reduce RF heating also reduce image artifact, because induced currents that cause tissue heating are the same responsible for producing secondary magnetic fields that distort scanner’s B1 field. To test this hypothesis, we performed experiments with a commercial DBS lead implanted in a human cadaveric brain undergoing MRI at 3T, with the extracranial portion of the lead configured along trajectories that produced maximum and minimum RF heating. We found that low-heating trajectories reduced image artifact around electrode contacts by up to 88% compared to high-heating trajectories.

Finally, we assessed the feasibility of implementing optimal lead trajectories in patients undergoing DBS surgery. We show that modified lead trajectories can be implemented in patients with minimal interruption in surgical workflow as described in case of a patient operated for subthalamic nucleus DBS in our institution.

## Materials and Methods

### Design and construction of anthropomorphic phantoms

We designed and constructed anthropomorphic phantoms based on computerized tomography (CT) images of a patient with implanted deep brain stimulation system. CT images of the patient were segmented into bone and soft tissue in 3D slicer(26) (Slicer 4.10, http://slicer.org) and output masks were further processed in a CAD tool (Rhino 6.0, Robert McNeal & associates, Seattle, WI) to create skull and torso-shaped structures. The skull structure consisted of two separable coronal halves, so that it could be re-filled with tissue-mimicking gel with different electrical prosperities. All parts were 3D printed in acrylonitrile butadiene styrene (ABS plastic) and coated with acrylic for waterproofing. Additionally, we designed and 3D printed grids and stands for supporting DBS leads extensions and IPG in locations analogous to clinical practice. The process of segmentation, 3D modeling, and construction of the phantom is illustrated in Figure 1.

**Figure 1:**
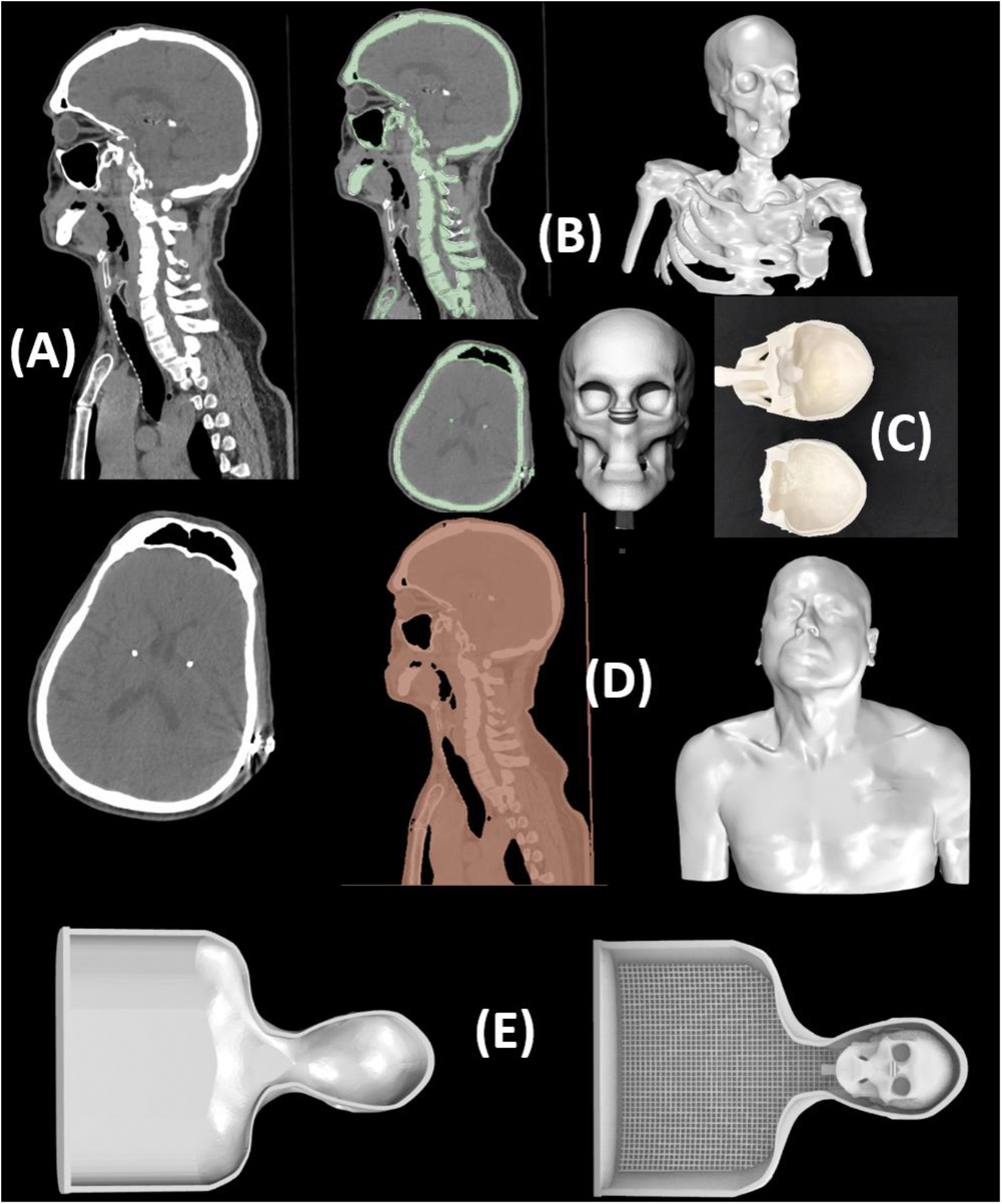
Design and construction of anthropomorphic phantom from CT images. (A) CT images of a patient with bilateral DBS implants. (B-C) Segmentation and creation of the bone structure from the CT images. (D-E) Segmentation and creation of body silhouette and fitted grids from the CT images.

### DBS device configurations

Two commercially available DBS systems with comparable lengths of leads and extensions were used in experiments: a Medtronic (Medtronic Inc, Minneapolis, MN) system and an Abbott (St. Jude Medical, Plano, TX) system including a 40 cm lead (Medtronic model 3387, Abbott model 6173), a 60 cm extension (Medtronic model 3708660, Abbott model 6372), and an IPG (Medtronic Activa PC-37601, Abbott Infinity, 6660). MR-compatible fluoroptic temperature probes (OSENSA, BC, Canada) were secured at electrode contacts 0 and 2 of the Medtronic lead and at electrode contacts 0 and 2b of the Abbott lead as illustrated in Figure 2. The lead-probe system was then inserted into the skull through a 5 mm hole, following entry point, angle, and penetration depth analogous to the clinical approach for targeting right subthalamic nucleus.

**Figure 2:**
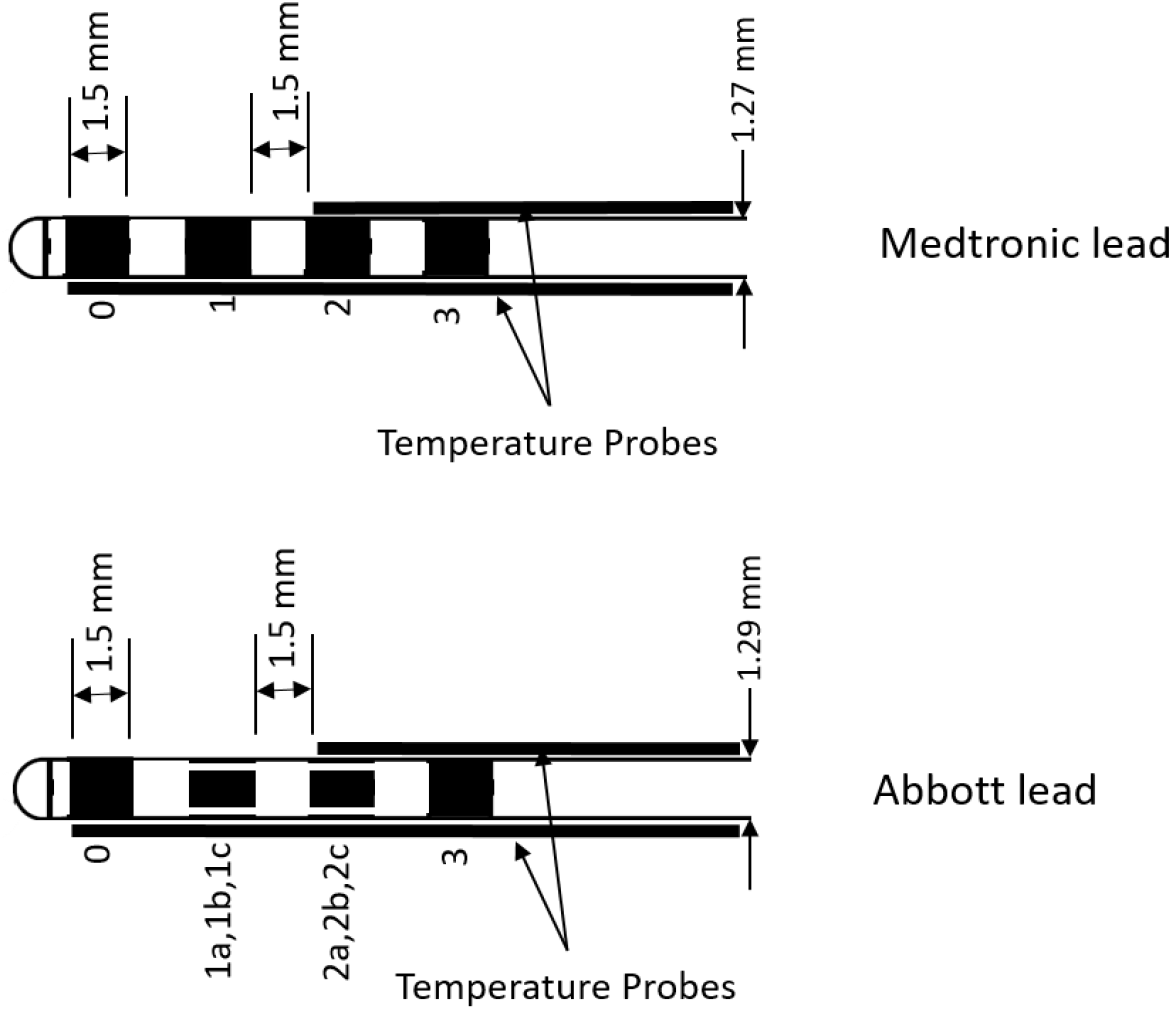
Schematic figures showing characteristics of the DBS leads and temperature probe positioning used in the experimental measurements.

**Figure 3:**
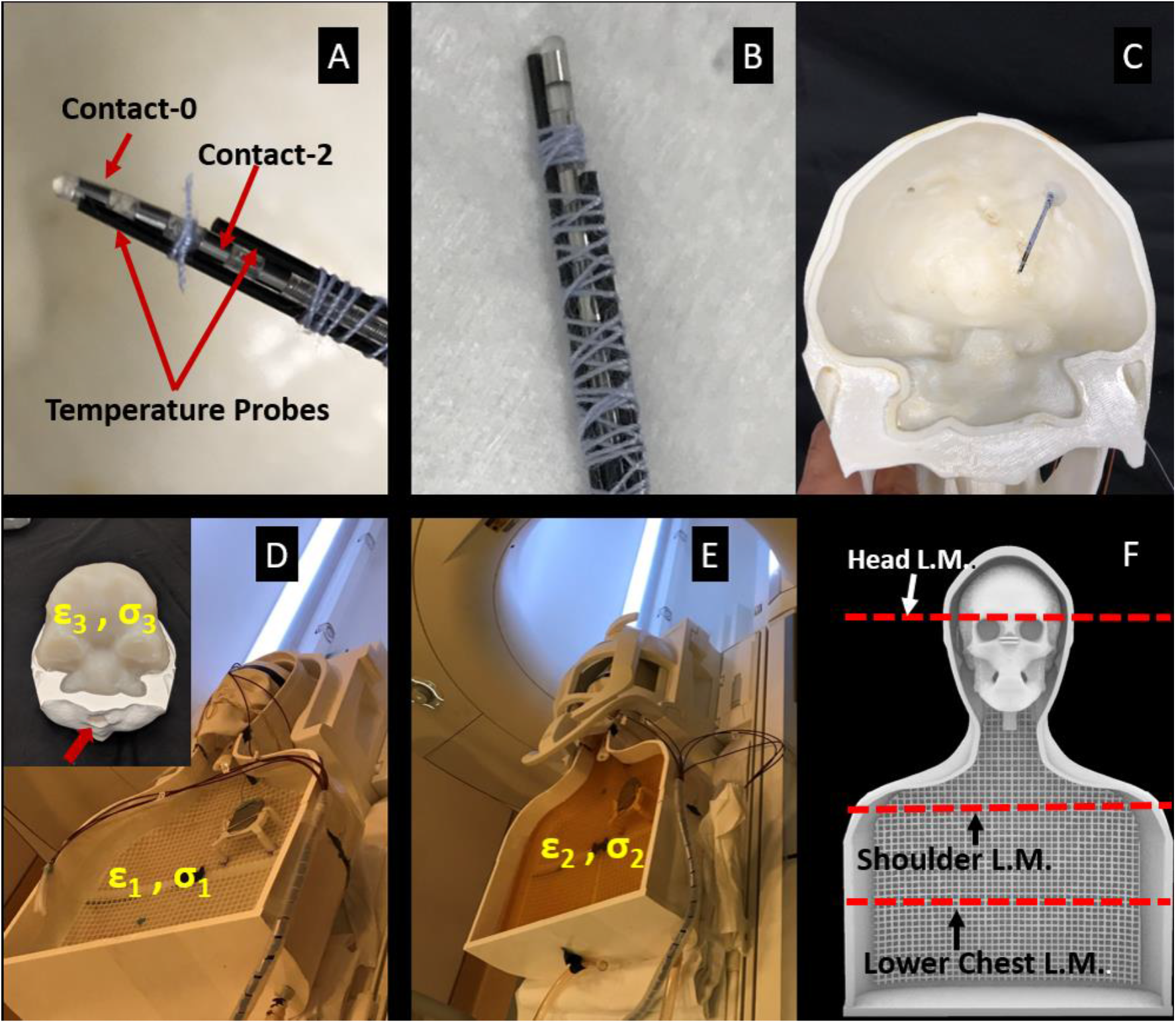
(A-B) Temperature probes secured at DBS lead contacts of Medtronic and Abbott DBS leads. (C) DBS lead attached to the temperature probes positioned into the skull. (D) Saline filled phantom implanted with the full DSB system. The inset shows agar gel inside the skull and a hole (red arrow) drilled on the base of the skull to allow conductive connection between the brain and the extracranial medium (saline). (E) Setup with additional oil layer. (F) Location of different imaging landmarks used for the heating measurements.

Tissue mimicking gel was prepared by mixing 32 g/L of edible agar (Landor Trading Company, gel strength 900 g/cm^2^) with saline solution (2.25 g_NaCL_/L) while heating with continuous stirring until a uniform solution was formed. The solution was let to cool down until its temperature fell below 60 °C, and then poured into the skull and let to further cool down to solidify. The conductivity and relative permittivity of the solidified gel was measured to be *σ* = 0.40 *S/m* and *ε_r_* = 78.

The gel-filled skull containing the lead-probe system was inserted into the head portion of the phantom. The lead was connected to the extension and IPG, with IPG positioned at the left pectoral region contralateral to the lead. This contralateral lead-IPG configuration has shown to systematically generate higher RF heating due to coupling of MRI electric fields with the medio-lateral tunneling segment of the lead (23).

For each DBS system, heating measurements were performed for eight lead-extension trajectories as shown in Figure 4. For trajectories 1 and 5, the lead was routed either mediolaterally toward the mastoid bone or in the anteroposterior direction toward the occipital bone without creating any loop on the skull. Positioning concentric loops in the extra cranial trajectory of DBS leads has shown to reduce RF heating in previous works (27,28), however, the optimal location of the loops has not been investigated. For all cases, the extra length of the extension was looped around the IPG as recommended by manufacturers.

**Figure 4:**
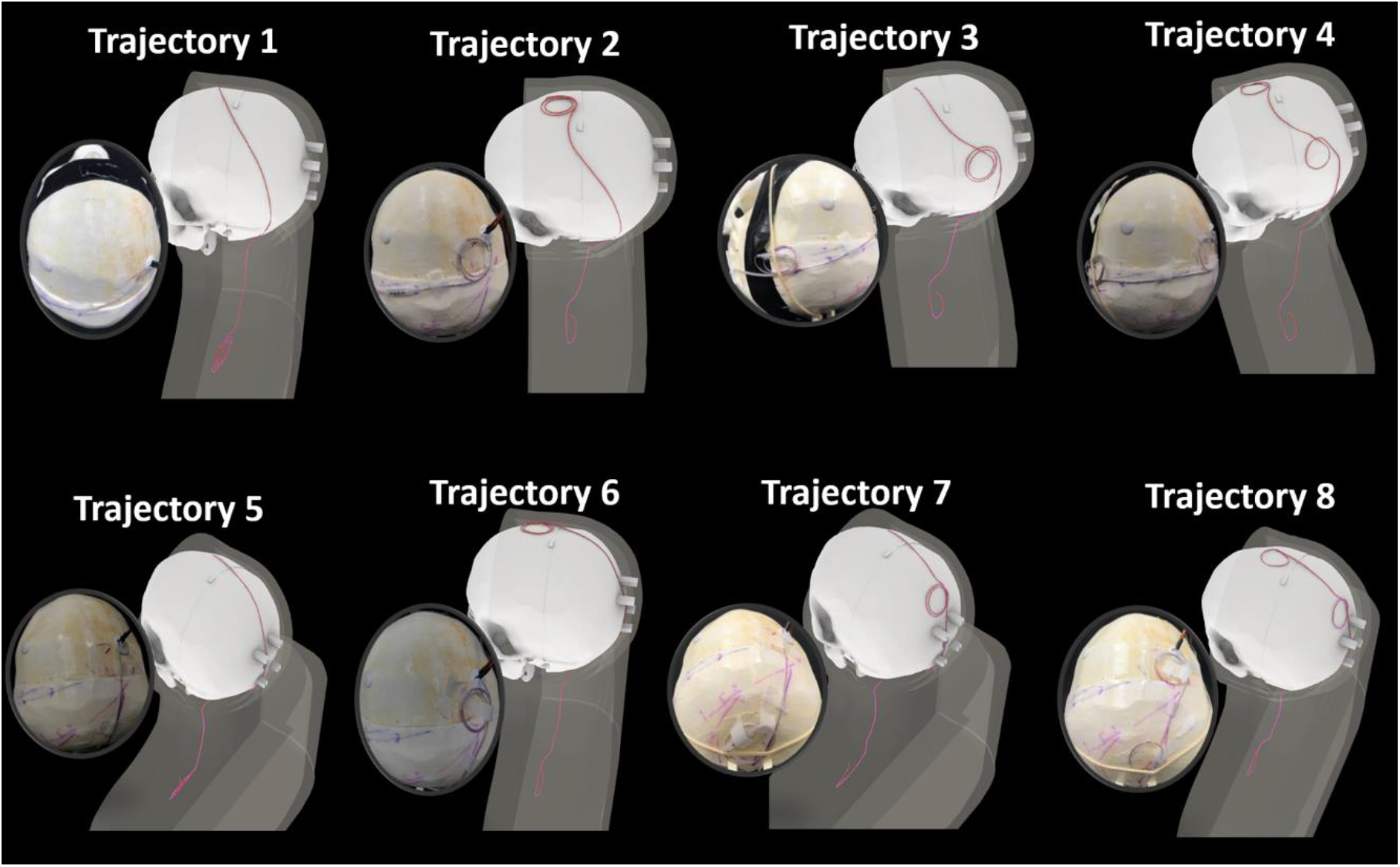
Different trajectories of the DBS lead/extension inside the anthropomorphic phantom.

To investigate the effect of body composition on the RF heating, each experiment was performed twice: one with the torso filled with saline (*σ* = 0.50 *S/m* and *ε_r_* = 78), and next with a thin layer of oil (*σ* = 0 *S/m* and *ε_r_* = 3) added on top of the saline solution representing subcutaneous fat. For the saline-only cases, we used 18 L of solution so the IPG as well as leads and extensions were fully submerged in the solution. For the cases with saline+oil, we replaced 7 L of saline with oil, such that the bottom layer of oil was touching the upper face of the IPG while other faces of the IPG were in contact with the saline.

### System validation and integrity verification

Before starting experiment for each device, we verified the integrity of implanted DBS system by measuring the inter-electrode as well as electrode-IPG impedances as recommended by the manufacturers. Measured impedance values were well within the manufacturer recommended range (< 4000 Ω for enter-electrode impedance and < 2000 Ω for electrode-IPG impedance). For Medtronic system, the IPG was set to ‘stimulation off’ mode and for Abbott system the device was set to ‘MRI mode’.

### RF exposure

Experiments were performed in a 1.5T Siemens Aera scanner and a 3 T Siemens Prisma scanner (Siemens Healthineers, Eglangen, Germany) using body transmit coil and a 20 channel head receive coil. A T1 weighted turbo spin echo (T1-TSE) sequence was used for both 1.5T and 3T experiments, with parameters adjusted to reach the maximum partial SAR limit as reported by the scanner console. Table 1 gives details of sequence parameters.

**Table 1:** Parameters for the sequences used for RF heating measurements at 1.5T and 3T.

| Sequence Parameters | 1.5 T | 3 T |
| --- | --- | --- |
| TE (mS) | 7.3 | 7.5 |
| TR (mS) | 814 | 1450 |
| Flip Angle | 150 | 150 |
| $B_1^{+rms}$ (μT) | 4.1 | 2.8 |
| Acquisition time (min) | 4:14 | 7:31 |

### Cadaver experiments

To study the image artifact around DBS electrodes for different extracranial lead trajectories, we used a cadaveric brain prepared and fixed in formalin positioned inside the open skull before the two halves were glued together. A Leksell model G base ring (Elekta, Stockholm, Sweden) was secured to the skull and the system was scanned at 1.5T along with the MRI fiducial box. The images were transferred to BrainLab iPlan server to determine the target and entry point using combination of standard stereotaxy formula coordinates for subthalamic nucleus and direct visualization of MRI images. An isolated Medtronic DBS lead (model 3387) was implanted into the subthalamic nucleus of the cadaver brain with the help of an arc attached to the base ring and a DBS cannula inserted 10mm proximal to the target. The skull was then placed into the anthropomorphic container filled with saline (*σ* = 0.48 *S/m*, *ε_r_* = 78) and imaged in a Siemens 3T Prisma scanner with the extracranial portion of the lead routed along trajectories 1 or 2.

## Results

### RF heating

Figure 5 shows the temperature rise,Δ*T*, at the most distal contact (contact-0) of the Medtronic lead for different trajectories, phantom compositions, imaging landmarks, and RF frequencies. RF heating was most severe during head imaging with Δ*Tmax* = 12.53°C at 1.5T and Δ*Tmax* = 4.72°C at 3T, followed by shoulder imaging with Δ*Tmax* = 3.70°C at 1.5T and Δ*Tmax* = 2.57°C at 3T, and lower chest imaging with Δ*Tmax* = 1.69°C at 1.5T and Δ*Tmax* = 0.61°C at 3T. Changes in DBS device configuration and patient’s body composition had substantial effect on the RF heating. Specifically, the temperature rise during head imaging increased from 3.75 °C to 12.53 °C at 1.5T and from 0.61 °C to 4.72 °C at 3T for trajectory 1 when a thin layer of oil was added on top of the saline solution in the torso, mimicking patients with more subcutaneous fat.

**Figure 5:**
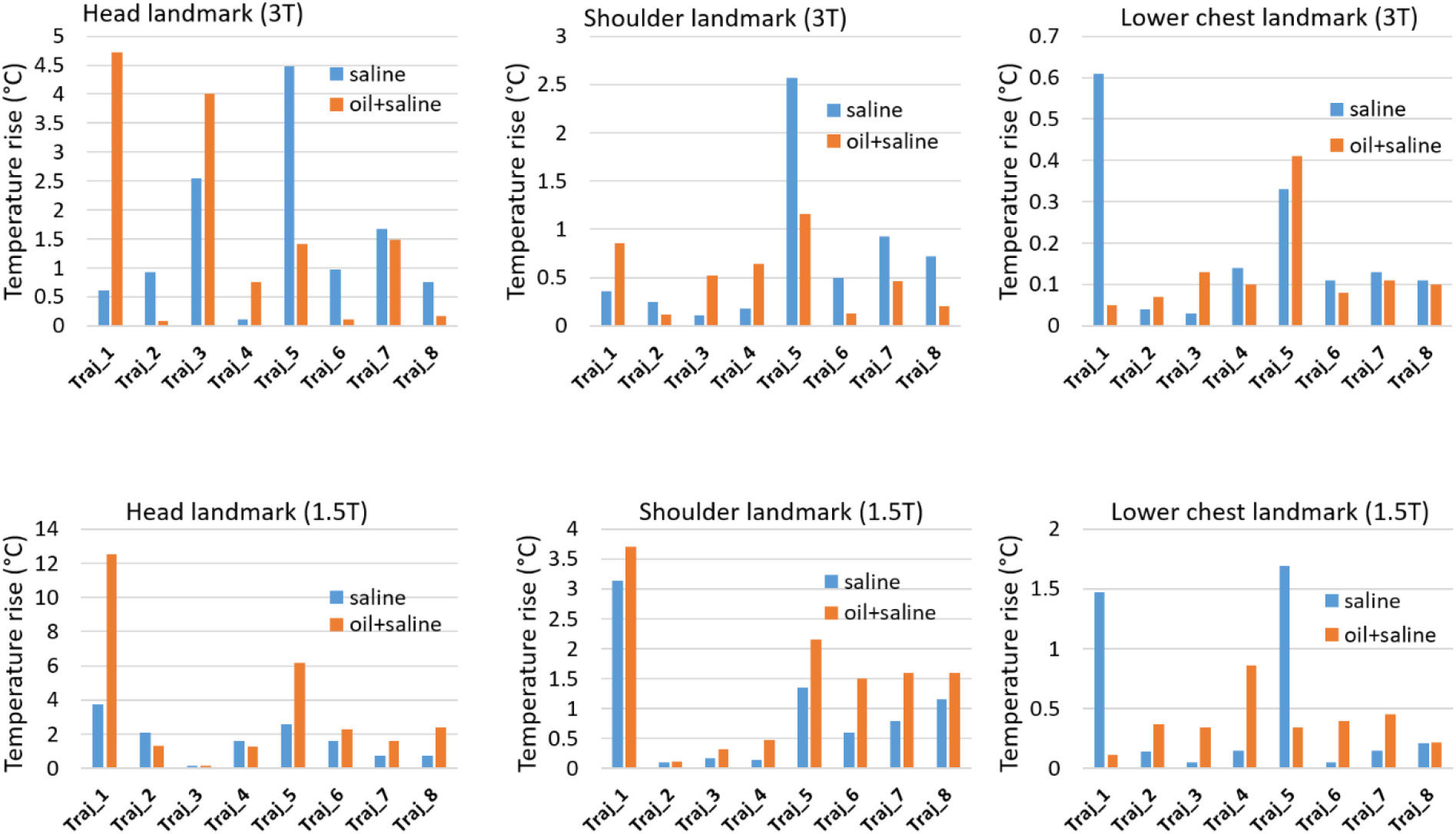
Plot of temperature rise for different imaging landmarks and body compositions for Medtronic DBS lead during MRI at 1.5 T and 3 T.

Figure 6 shows the temperature rise,ΔT, at the most distal contact (contact-0) of the Abbott DBS device for different trajectories, phantom composition, imaging landmark, and RF frequencies. Similarly, maximum temperature increase was observed during head imaging, Δ*Tmax*=23.73°C at 1.5T and Δ*Tmax*=6.60°C at 3T), followed by shoulder imaging (Δ*Tmax*=7.97°C at 1.5T and Δ*Tmax=7.39°C* at 3T), and lower chest imaging (Δ*Tmax*=2.22°C at 1.5T and Δ*Tmax*=1.87°C at 3T). The effect of variation in body composition was most prominent for trajectory 1 at 1.5 T with Δ*T* increasing from 4.68°C to 23.73°C when a thin layer of oil was added on top of saline solution in the torso phantom.

Figure 7 gives the mean and standard deviation of RF heating for each lead trajectory, averaged over different phantom compositions, imaging landmarks and MRI RF frequencies (64 MHz at 1.5T and 127 MHz at 3T).

**Figure 6:**
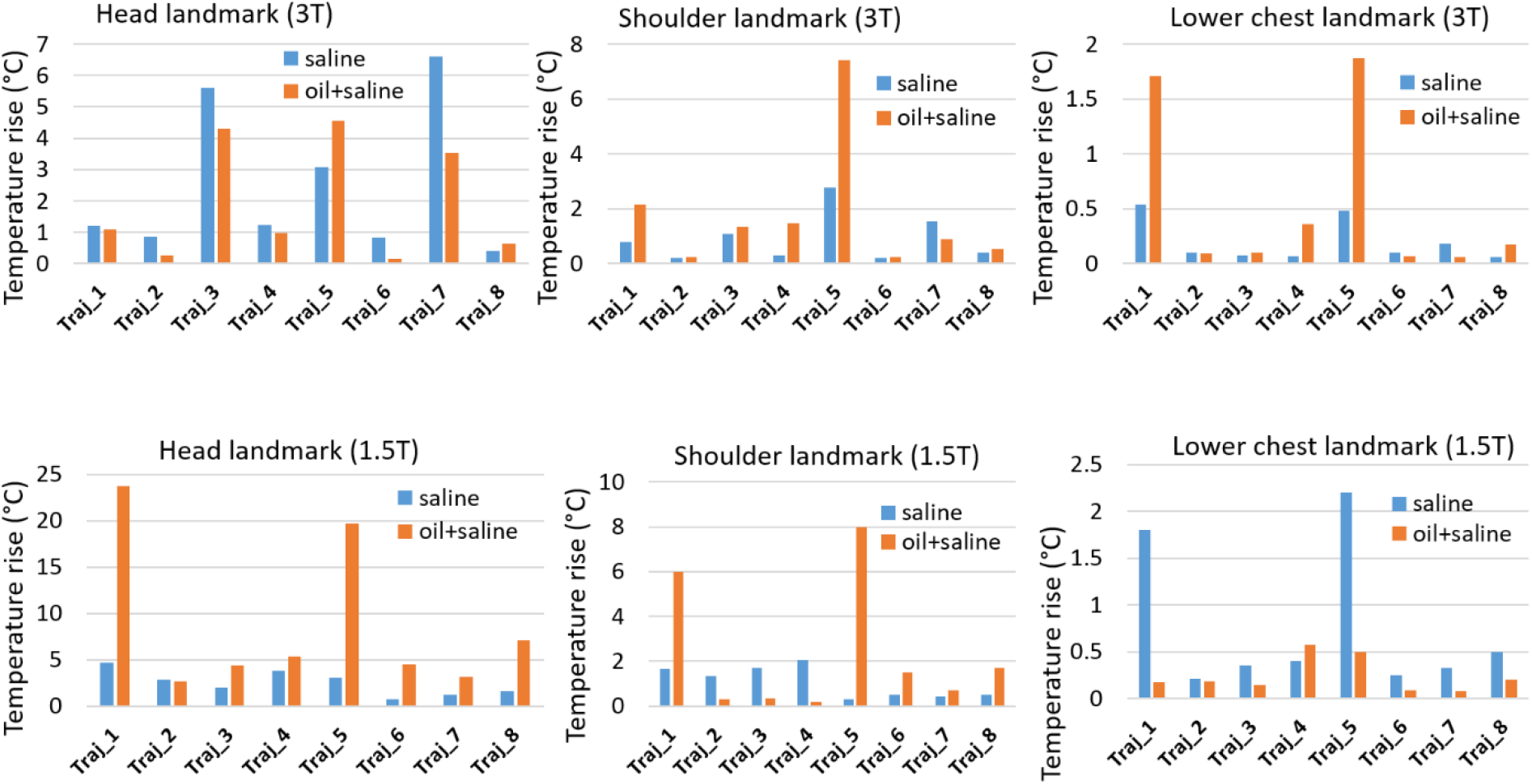
Plot of temperature rise for different imaging landmarks and body compositions for Abbott DBS lead during MRI at 1.5 T and 3 T.

**Figure 7:**
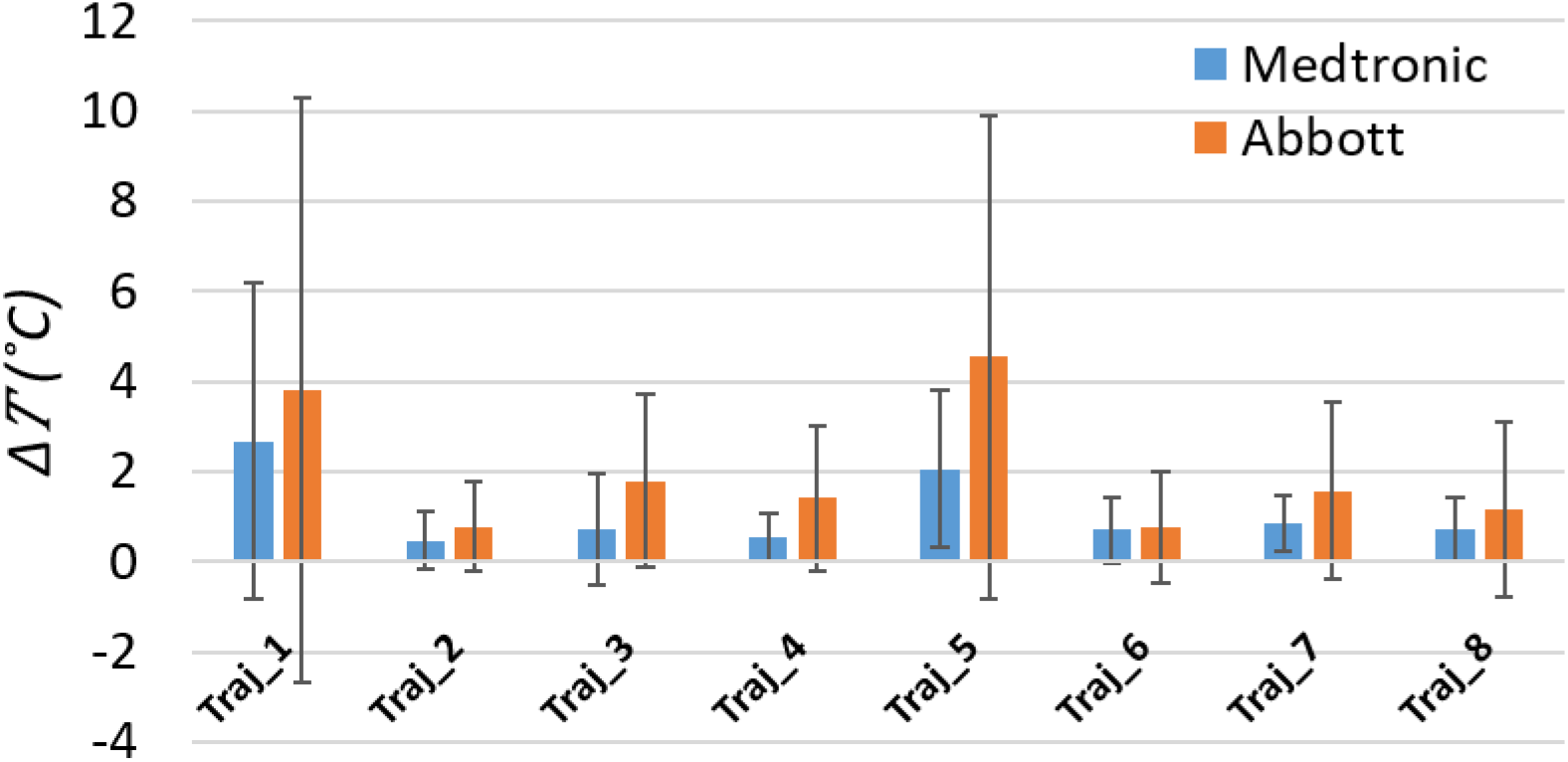
Plot of mean and standard deviation of the temperature rise for each device for individual trajectory, averaged over all imaging landmarks, Scanner field strengths and body compositions.

Considering all the variants of the study, minimum heating occurred for the trajectory with concentric loops at the surgical burr hole with the lead routed medio-laterally towards the ear and the excess length of extension looped about the IPG (trajectory 2).

### Image artifact

Figure 8 shows the image artifact around the DBS lead in the cadaver brain in coronal and transverse planes, for extracranial lead trajectories associated with maximum (A, C) and minimum (B,D) RF heating. Images were imported into MATLAB (MathWorks Inc., Natick, MA), where the artifact around the lead was manually masked out and the number of affected pixels were counted. Compared to the straight lead trajectory, optimal trajectory with concentric loops reduced image artifact by 88% in the transverse plane and by 50% in the coronal plane.

**Figure 8:**
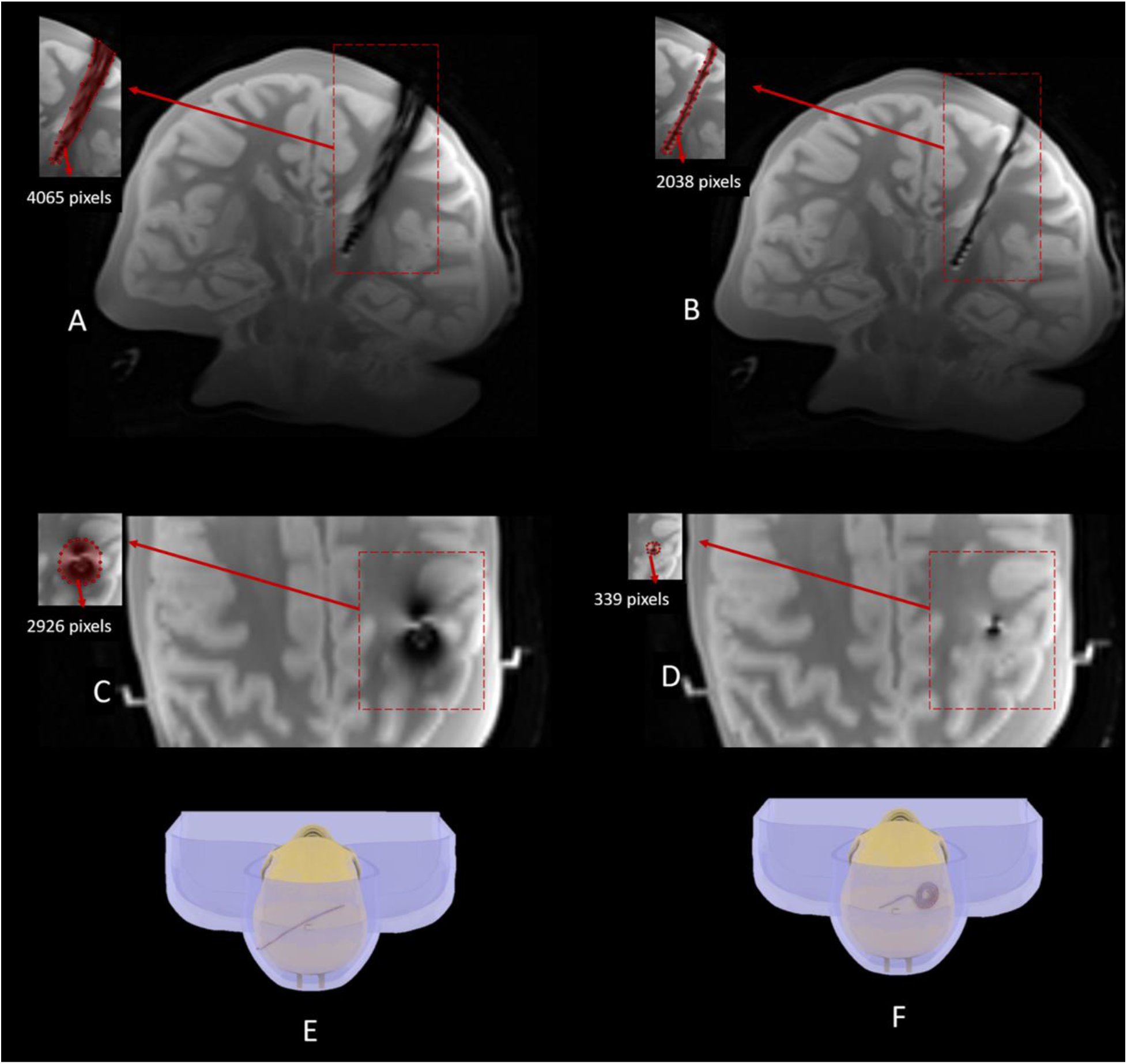
Image artifact in coronal and transverse planes in a cadaver brain implanted with DBS leads for two different trajectories of extracranial lead: no loop on the skull (E) and concentric loops at the burr hole (F).

### Surgical implementation

From the practical perspective, optimal trajectories can be implemented in patients with minimum disruption to the surgical flow. One example is given in Figure 9 showing postoperative CT images of a patient receiving bilateral DBS implants targeting subthalamic nucleus at one of our institutions (Albany Medical Center). For guiding the lead trajectory, we used curved mayo scissors passing posterior and to the left of the incision. The blades of the scissors were then opened to their widest to create a pocket for placing loops of each lead. The leads were looped in two to three concentric circles which were then placed at the lead pockets. The rest of the leads were passed toward the temporal lobe where they would be connected to their respective extensions. The extensions were tunneled towards left chest region and connected to a dual channel IPG implanted in left pectoralis, as shown in Figure 9.

**Figure 9:**
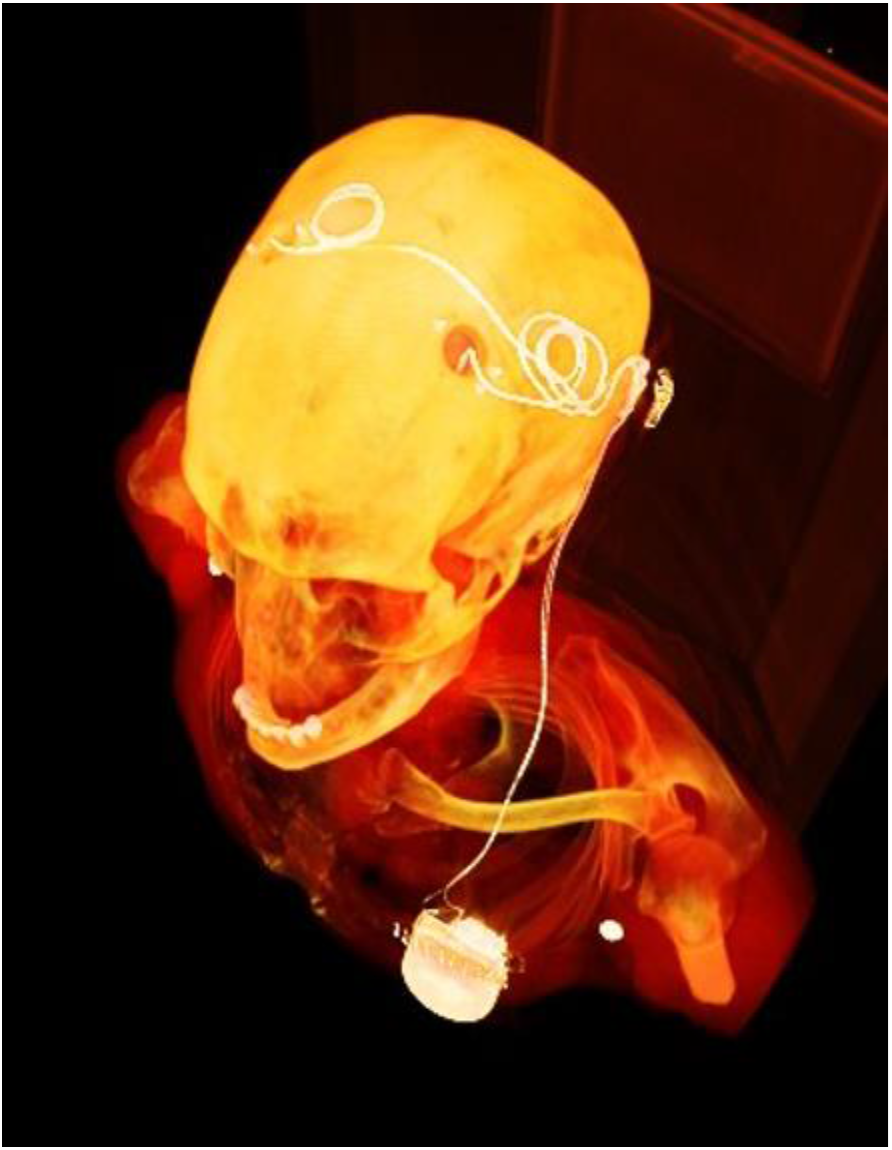
CT Images of a 62 years old male with Parkinson’s disease receiving bilateral DBS implants following the optimal device trajectory for the reduction of RF induced heating and image artifact during MRI.

## Discussions and Conclusions

In recent years, there have been increased efforts to mitigate the problem of RF heating of DBS as well as other type of elongated metallic implants during MRI. Some efforts aim to change the MRI hardware to reduce the interaction of transmit electromagnetic fields from the MRI scanner with the implanted leads(15,18,19,29–36). Others, have purposed the modification of lead design and the materials to suppress the induced currents produced by the lead interacting with the transmit fields from the scanner(37–39). One low cost and easy to implement strategy to reduce RF heating is modification of trajectories of the DBS leads and extensions. Different mechanisms are proposed by which concentric loops could reduce RF heating of DBS leads. One is that looping the lead will create additional inductance, increasing the overall impedance of the lead and thus reducing induced RF currents. Another possibility is that the change of magnetic flux in the loop during B1 excitation creates electromotive forces and currents which counteract RF currents induced by MRI electric fields (28). It has been also suggested that the reduction of RF heating is the direct result of MRI electric fields being cancelled on the opposite sides of the loop (23). Although the SAR-reduction effect of loops has been shown in both simulations and measurements, the optimum positioning of the loops for maximum SAR reduction was not systematically evaluated. Theoretically, because RF heating is a resonance phenomenon, the optimum position of the loop will depend on the length of the lead/extension in hand. This work presented a systematic study of effect of different loop positioning for two commercially available DBS devices with comparable length of lead and extension (40cm and 60 cm, respectively), considering different phantom compositions, imaging landmarks, and RF frequencies. Our results show that introducing concentric loops at the location of surgical burr hole consistently reduces the RF heating. Particularly, this trajectory reduced RF heating of the Medtronic and Abbott devices used in this study during head MRI by 89% and 88% at 1.5T and by 98% and 87% at 3T respectively, compared to corresponding worst heating scenario. However, as the length of leads and extensions varies considerably between different models (30-40 cm for the lead, and 40-90 cm for the extension) more work is required to assess optimum trajectories for other combinations of lead and extensions. Hypothetically, such study may find that a universal trajectory is optimum across all body types, sex, and device models, or give recommendation in the form of “For a male patient receiving a Medtronic implant, implement trajectory A”, and so on.

Another important factor that has been overlooked is the effect of patient’s body composition on RF heating of DBS implants. As dielectric properties of fatty tissue are significantly different from those of muscular tissue or the average tissue properties, the electromagnetic phenomena in a multi-compartment sample could lead to a different heating pattern than what is observed in a single-material phantom.

Most of the previously reported studies assessing RF heating of DBS implants have used either homogenous body models or single material gel phantoms. The work presented here is the first to highlight the effect of patient’s body composition, especially the subcutaneous fat around the DBS device, on RF heating at electrode contacts. The presence of fat substantially altered RF heating and up to 7-fold increase in heating was observed in some cases. Our results suggest that studies that aim to evaluate MRI safety in patients with DBS implants should also consider the effect of differences in dielectric properties of tissues surrounding the implanted device.

Another important issue is the image artifact around DBS leads. Metal introduces both a susceptibility artifact due to B_0_ distortion, and an RF artifact due to B_1_ distortion from secondary magnetic fields generated by induced currents on the implant. The latter is particularly strong in wire implants and has been used to characterize induced currents (40). As most modern implants are made of non-ferromagnetic material with susceptibility close to water, the RF artifact is the dominant factor degrading image quality. Theoretically, a low-SAR trajectory also reduces RF artifact as the currents that cause RF heating are the same as those that cause local distortion in the B1 field. In this work we showed that such RF artifact could have a detrimental effect on the image, and that it can be substantially reduced by trajectory modification.

Finally, because optimizing lead trajectories is only useful to the extent they are implemented by surgeons, complimentary work is required to develop surgical techniques that ensure optimal trajectories require minimum disruption to the surgical flow. Here we show that such technique could be implemented in patients with minimal disruption to the surgical flow. In our case, the surgery was completed without complications, with trajectory modification adding only ~10 minutes to the typical 4-hour surgery. From manufacturer point of view, altering lead’s design and material to reduce RF heating has a substantially high cost-to-benefit ratio as the safety margins gained are modest at best, whereas slight alterations in design or material trigger the long and costly process of re-obtaining approval from regulatory agencies, for biocompatibility, non-toxicity, and mechanical stability. On the other hand, modifying MRI hardware and improving intra-surgical lead management have the potential to dramatically improve safety margins in ways that are generally applicable to a wide range of devices, thus dramatically shifting the cost-benefit calculation. Notably, as the network of high-volume DBS surgeons in the US remains small (approximately 100 implant centers), establishing such changes has been widely successful. For example, surgical practice quickly adapted once it became clear that the extension connector should be placed above the mastoid bone to avoid lead fracture – a surgeon-initiated modification that led to a significant decline of device complications. Similarly, innovative strategies to leverage surgical investment in minimizing SAR through optimized routing of extracranial DBS leads/extensions has the potential to become widely adopted, thus positively impacting patient’s care in large scale.

## Notes

**Grant Support** This work was supported by NIH funding with grants R00EB021320 and R03EB025344.

### Competing Interest Statement

The authors have declared no competing interest.

